# Phenotypic screens identify biologic regulators of nanoparticle uptake in diffuse midline glioma

**DOI:** 10.64898/2026.09.08.749175

**Authors:** Emmeline L. Cheng, Sangita Pal, Jessica W. Tsai, Clara B. Yampanis, Shriya Rangaswamy, Julianna Kenny-Serrano, Marissa Coppola, Merve Ozdemir, Kelly Y. Cai, Ethan VanGosen, Emma Bateman, Kimberly R. Bennett, Margaret M. Billingsley, Charles A. Whittaker, Ye Zhang, Myra Dada, Cong Fu, Sonam Bhatia, Yeji Bae, Neerja Katiyar, John G. Doench, Keith L. Ligon, Scott R. Manalis, Paula T. Hammond, Pratiti Bandopadhayay, Joelle P. Straehla

## Abstract

Nanoparticle drug delivery systems hold considerable promise for locoregional administration to central nervous system tumors, yet the biological determinants of nanoparticle-cancer cell interactions remain poorly understood. Using patient-derived histone-mutant diffuse midline glioma (DMG) models, we performed a pooled CRISPR-Cas9 perturbation screen to systematically identify regulators of liposomal nanoparticle delivery. The screen identified candidate genes spanning endocytosis, vesicle transport, and metabolic signaling, revealing that nanoparticle delivery is governed by a broader landscape than previously appreciated. Among these, *CTNNB1*, or β-catenin, emerged as a common negative regulator across two independent DMG models and two distinct nanoparticle surface chemistries. Transcriptomic profiling of *CTNNB1*-depleted DMG cells revealed upregulation of membrane remodeling and extracellular matrix gene programs, accompanied by reduced cell stiffness measured by a microfluidic acoustic scattering assay. This resulted in a shift in endocytic activity characterized by decreased bulk-phase macropinocytosis and increased receptor-mediated endocytosis. We further identified MAPK and mTOR pathway members as nanoparticle trafficking modulators, and demonstrated concordance between genetic and pharmacologic perturbations in modulating the liposomal nanoparticle interactions in pediatric DMG cells. These findings establish a biology-first screening approach for identifying previously unappreciated regulators with potential relevance to nanoparticle-based therapeutic strategies in pediatric brain tumors.

## INTRODUCTION

Nanoparticle drug delivery systems have emerged as a versatile platform for controlled delivery of therapeutic cargos, with a particular promise for locoregional administration to central nervous system tumors where direct access circumvents the blood-brain barrier and enables high local concentration at the target site^1^. Despite this potential, the biological determinants of successful nanoparticle-cancer cell interactions remain poorly identified, representing a fundamental barrier to nanoparticle design^2^.

Pediatric diffuse midline glioma (DMG) exemplifies this unmet clinical need. Pediatric DMG is a devastating subset of glial tumors that are nearly universally fatal. These tumors, driven by recurrent mutations in the histone protein H3 arise in midline structures such as the pons or thalamus that preclude surgical resection^3–5^. Radiation remains the standard of care but only provides transient benefit extending median survival from 3-6 months to 9-12 months^6^. Drug-delivery to the pons is a major challenge in the design of new therapeutic approaches for children with DMGs. Recent phase I studies have established the safety of catheter-directed locoregional therapy, delivering small molecules, cellular therapies, and liposomal therapeutics directly to the tumor or cerebroventricular system^7–9^. To expand the potential of nanoparticle therapeutics for locoregional administration, we sought to interrogate the biologic regulators of nanoparticle delivery to pediatric DMG cells, with the goal of identifying strategies to enhance nanoparticle uptake.

Nanoparticle delivery is known to depend on physicochemical parameters including particle shape, surface chemistry and core stiffness^10–12^, but comparatively little is known about how the biological state of target cells regulates uptake. Pooled screening approaches offer an efficient strategy for interrogating these biological interactions at scale^13^. Fluorescence-activated cell sorting (FACS) has recently been combined with nanoparticle screening and applied across cancer cell panels^14^ and genetically perturbed cells^15,16^, demonstrating that unbiased screens can uncover individual genes and pathways important for delivery that would not emerge from hypothesis-driven approaches. However, such screens have not been applied to pediatric brain tumor models, and the regulators governing nanoparticle uptake in this context remain unknown.

Here we describe a pooled CRISPR-Cas9 perturbation screen designed to systematically identify positive and negative regulators of liposomal nanoparticle delivery in patient-derived histone mutant DMG models. Liposomal nanoparticles were selected given their established clinical utility and direct translational relevance to locoregional delivery in pediatric DMG. This work establishes a biology-first screening paradigm for interrogating nano-bio interactions and identifies β-catenin and MAPK signaling as previously unappreciated regulators of nanoparticle trafficking in pediatric glioma.

## RESULTS AND DISCUSSION

### Pooled CRISPR-Cas9 screening identifies functional regulators of nanoparticle association in DMG

To identify the genetic determinants of nanoparticle-cell interactions in DMG, we performed a functional knockout screen using two patient-derived neurosphere models (BT245 and BT869) (Fig. 1a). This perturbation-based approach allows for identification of genes that causally regulate nanoparticle association and/or trafficking. We utilized a library of 3,120 sgRNAs targeting 705 genes associated with endocytosis, vesicle transport, and metabolic signaling, including nodes previously implicated in nanoparticle association across diverse cancer contexts^14^ (Fig. 1b).

To evaluate the influence of surface chemistry, we used layer-by-layer assembly to generate two nanoparticles with identical fluorescent liposomal cores but distinct surface properties. Hyaluronic acid (HA) was chosen for the outer layer for this screen due to high affinity for glioma models and potential for enhanced delivery across the blood-brain barrier^17,18^. Nanoparticles size, polydispersity (PDI), and surface charge were characterized prior to each experiment and confirmed successful layer-by-layer assembly through expected shifts in surface zeta potential (Fig. 1c). Both formulations maintained a hydrodynamic diameter <130nm and uniform distribution with polydispersity index < 0.2.

**Fig. 1.**
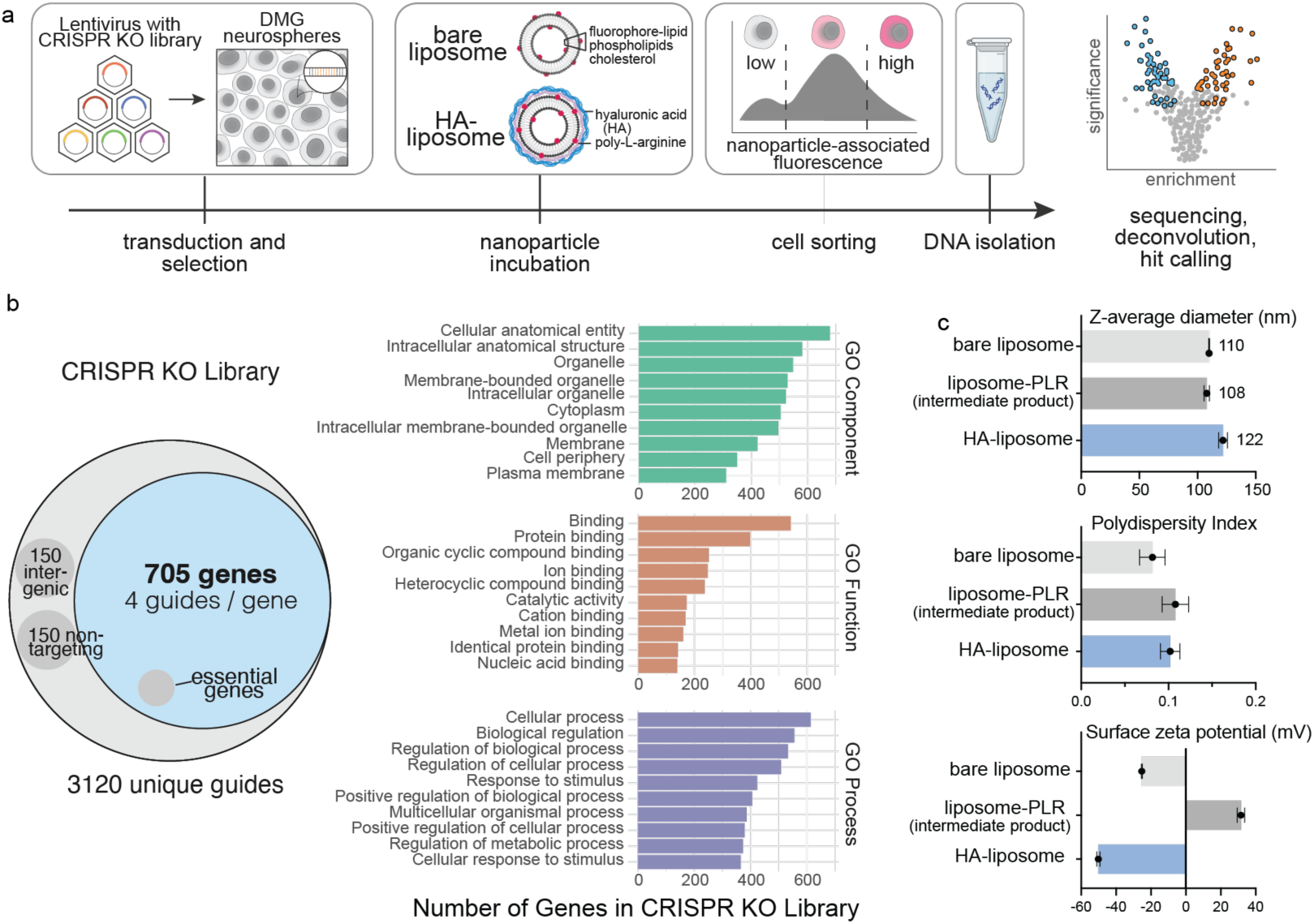
Schematic of experimental workflow and screen details. **a**, Schematic showing screen workflow. Diffuse midline glioma (DMG) neurospheres were infected with a CRISPR knockout (KO) library, followed by incubations with bare liposome and HA-liposome variations, after which flow sorting was done based on degree of liposome-associated fluorescence. High and low fluorescently labeled cell populations from top and bottom quartiles were sequenced for guide enrichment and hit calling. **b**, Gene ontologies of the CRISPR KO library encompassing 705 target genes. **c**, Nanoparticle characterization of the bare liposome and the HA-liposome used for the screen.

Following lentiviral transduction and antibiotic selection of the library^19^, DMG neurospheres were incubated with fluorescent nanoparticles for 1, 4, or 24 hours. Using fluorescence-activated cell sorting (FACS), we isolated the top and bottom quartiles of nanoparticle-associated cells, and measured sgRNA abundance within each of them. Robustness of dynamic gating was confirmed by consistent fluorescence distributions across technical replicates (Supplementary Fig. 1a-c). Quality control studies demonstrated high library coverage and representation throughout the assay (Supplementary Fig. 2). The top quartile includes cells with high fluorescence and therefore higher nanoparticle uptake, while the bottom quartile includes cells with low nanoparticle uptake. Comparisons in sgRNA abundance within these two groups allowed us to identify those that are specifically depleted in cells with high nanoparticle association. Genes targeted by each of these sgRNAs thus represent candidate targets whose ablation enhances nanoparticle trafficking (Fig. 2). Conversely, genes targeted by sgRNAs enriched in the bottom quartile represent positive regulators of nanoparticle uptake and therefore those whose ablation inhibits nanoparticle uptake.

We applied our nanoparticle CRISPR library to two patient-derived histone-mutant DMG neurosphere cell lines – BT245 (derived from a thalamic DMG) and BT869 (derived from a pontine DMG). Based on initial kinetics in our screen performed on BT245 that yielded consistent candidate gene number at 4- and 24-hour timepoints, we opted for shortened timepoints at 1- and 4-hour which did not impact hit count (Fig. 2a, Supplementary Fig. 3). Overall, our screen identified 55 and 36 high-confidence modulators of nanoparticle trafficking in BT245 and BT869, respectively (Supplementary Fig. 3), with both overlapping and diverged hits between the two models (Supplementary Fig. 4).

Analysis of guide enrichment across both DMG models identified β-catenin (*CTNNB1*) as a dominant and previously undescribed regulator of nanoparticle association (Fig. 2a). In the BT245 model, *CTNNB1* depletion was strikingly correlated with increased nanoparticle uptake at both the early and late timepoint (bare liposome FDR < 0.0001 for both early and late; HA-liposome FDR < 0.0001 for both early and late). In the BT869 model, *CTNNB1* depletion also correlated with increased nanoparticle uptake at the early timepoint (bare liposome FDR = 2 x 10^-3^, HA-liposome FDR < 0.0001) but was not significant at the late timepoint. Although β-catenin is a well-recognized scaffold for cell adhesion^20^ and transducer of Wnt signaling^21^, its role in modulating synthetic nanoparticle entry in the context of DMG has not been previously explored. Given the robust and cross-model nature of this hit, we prioritized *CTNNB1* for mechanistic characterization and functional validation in subsequent experiments, seeking to define its role in nanoparticle trafficking in DMG.

In contrast to *CTNNB1* which was scored across models and nanoparticle chemistries, we also identified targets that were model- and surface-chemistry dependent. Notably, exostosin glycosyltransferase 1 (*EXT1*) emerged as a positive regulator of nanoparticle uptake, with *EXT1* depletion significantly correlated with decreased nanoparticle uptake exclusively in the BT245 model and specifically in HA-liposomes (FDR < 0.0001 for both early and late timepoints, Fig. 2a). Whereas CTNNB1 depletion increased nanoparticle uptake globally, EXT1 depletion markedly reduced uptake of HA-liposomes without impact on bare liposomes. This specificity may be related to the polysaccharide coating on the liposomes, as *EXT1* is a glycosyltransferase that plays a key role in regulating proteoglycans on the cell surface^22^. The identification of *EXT1* highlight the potential for pooled, phenotypic screens to resolve model-specific dependencies; however, its relevance appears restricted to the BT245 model within the observed timeframes. It remains possible that *EXT1* could emerge as a regulator in BT869 at later timepoints. While providing valuable insight into the fundamental biology of HA-liposome delivery, the impact of *EXT1* depletion on nanoparticle delivery contrasts with our objective to enhance delivery efficiency and led us to prioritize targets with more direct translational potential.

Beyond individual outliers, pathway analysis of convergent hits identified many genes associated with activation of the mitogen-activated protein kinase (MAPK) and mechanistic target of rapamycin (MTOR) pathways (Fig. 2b). These hits clustered almost exclusively as negative regulators, wherein genetic depletion was correlated with increased nanoparticle association. The MAPK family transmits a wide range of extracellular stimuli to modulate gene expression, regulating crucial cellular functions^23^. Although the library was not specifically designed to tile these pathways comprehensively, we observed significant enrichment of key upstream regulators such as EGFR, FGFR1^23,24^, and PTPN11 (SHP2)^25,26^. We also identified the downstream multidrug resistance transporter ABCB1 (P-glycoprotein), which is known to be transcriptionally modulated by MAPK signaling^27,28^. The signaling signature was further complemented by the identification of parallel regulators including the PTEN/PTENP1 locus and MTOR itself. The emergence of these nodes is particularly noteworthy given their roles as either suppressors of the MAPK cascade^29,30^, or participants in compensatory feedback loops^31^.

This enrichment was often model-specific, a divergence likely underscored by the distinct genetic backgrounds of our models (Fig. 2c). For instance, PTEN emerged as a top negative regulator in BT869 but not in BT245, where PTEN is already non-functional due to a primary mutation, illustrating the screen’s ability to resolve hits within the context of existing oncogenic drivers. While these hits exhibited varying degrees of model-dependency, their shared association with extracellular stimulus processing and inherent pharmacologic tractability make them attractive targets for delivery modulation. Overall, our screen identified a preponderance of negative regulators over positive ones. This profile diverges from a previous nanoparticle screen using gold nanoparticles^15^ but aligns with another employing population-matched controls^32^, suggesting our short-term, targeted approach successfully prioritized specific modifiers of uptake while minimizing fitness-related artifacts.

Taken together, our phenotypic nanoparticle screen resolved both a singular, high-confidence regulator in *CTNNB1* and a group of broader, clinically actionable signaling pathways as regulators of nanoparticle delivery to DMG cells. Based on these findings, we focused our validation studies on the mechanistic role of *CTNNB1* in nanoparticle entry and potential for pharmacologic inhibition of the MAPK pathway to modulate delivery.

**Fig. 2.**
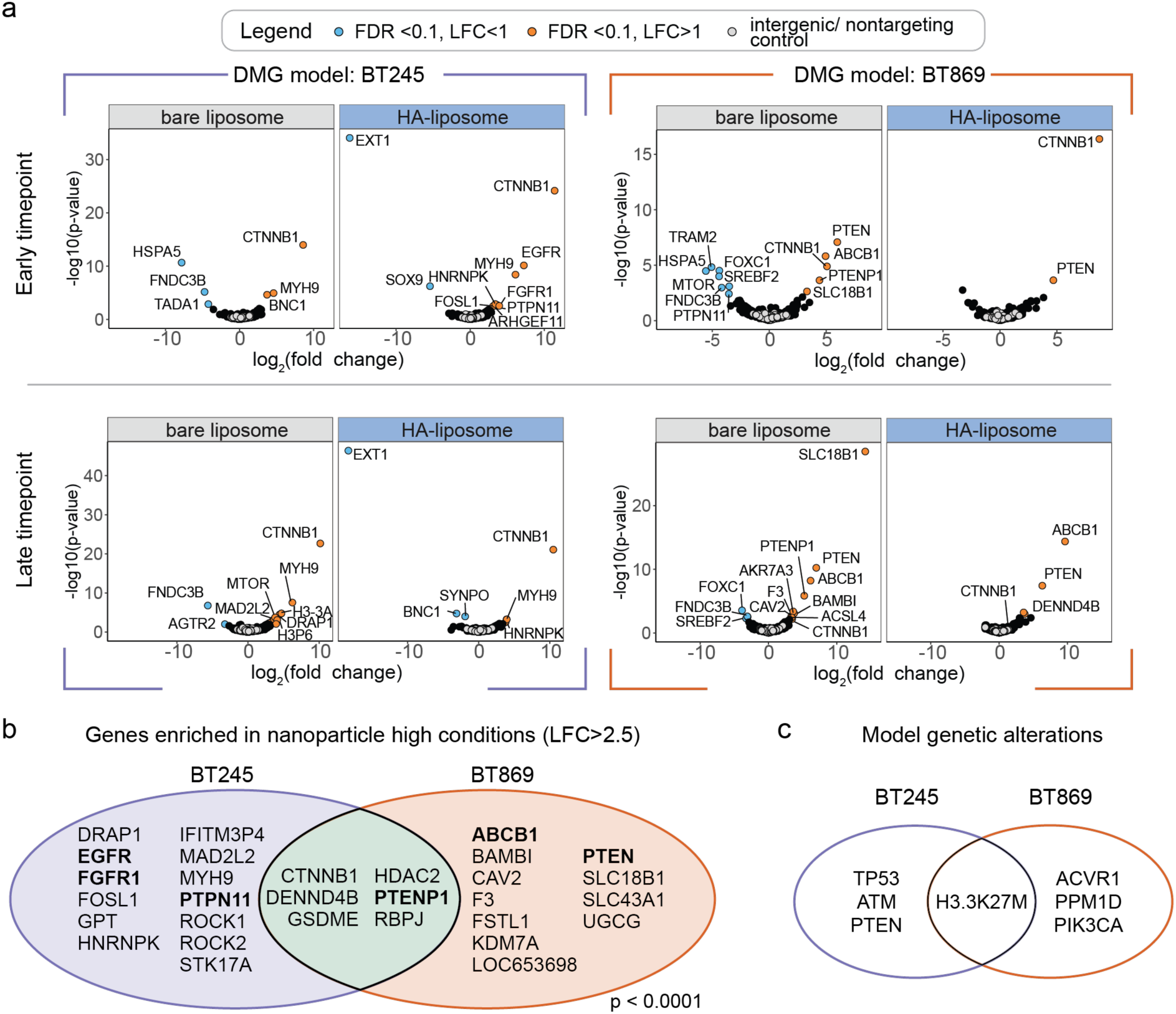
Screen results. **a**, Guide enrichment in BT245 (left) and BT869 (right) DMG neurosphere models after treatment with bare liposomes or HA-liposomes at early and late timepoints. X-axis represents fold-change abundances in guides in cells sorted in the top quartile (positive log-fold change) or in the bottom quartile of fluorescence (negative log-fold change). Y-axis depicts significance (-log10, *p*-value, the Broad Institute Apron analysis tool). **b**, Venn diagram of unique and overlapping top enriched genes in nanoparticle-high conditions (*p* < 0.0001, Fisher’s exact test). **c**, Venn diagram of major genetic alterations in BT245 and BT869 models.

### CTNNB1 knockout increases uptake of lipid and nonlipid nanoparticles in DMG models

To determine whether *CTNNB1* acts as a negative regulator of nanoparticle delivery, we generated isogenic *CTNNB1*-knockout models in both DMG cell lines (BT245 and BT869). Four sgRNAs targeting *CTNNB1* were included in the phenotypic screen, and the two best performing guides were selected for validation assays alongside a control non-targeting guide (sgGFP) (Supplementary Fig. 5a). We confirmed *CTNNB1* knockout via Western blot (Supplementary Fig. 5b-c) prior to proliferation assays. Quantification of intracellular ATP levels and neurosphere size confirmed that both DMG models tolerated *CTNNB1* loss without changes in cell fitness (Supplementary Fig. 5 d-g).

We next sought to validate our pooled screening result using an analogous workflow (Fig. 3a). As *CTNNB1* was a candidate regulator for both liposomal nanoparticles in the screen, we included a non-lipid nanoparticle (carboxylated polystyrene), as a control particle that was not used in the pooled screen. After selection, DMG cells were incubated with fluorescently labeled bare liposomes, HA-liposomes, or polystyrene nanoparticles, and nanoparticle association was measured by flow cytometry. Consistent with the pooled screen, *CTNNB1* depletion increased association with both bare and HA-liposomes compared to control cells. This relationship also extended to the nonlipid nanoparticle. All three nanoparticle types showed significantly higher association with *CTNNB1*-knockout cells over control cells (*p* < 0.01 or *p* < 0.001), with the largest absolute increases seen for HA-liposomes and nonlipid nanoparticles (Fig. 3b-c, Supplementary Fig. 6). The inverse relationship between *CTNNB1* expression and nanoparticle association was observed across technical and biological replicates (Fig. 3d).

Concordance between the screen and validation assays, along with replication across models and nanoparticles, supports a robust relationship between *CTNNB1* and nanoparticle association in DMG cells. The extension to non-lipid nanoparticles suggests the effect may be generalizable and as opposed to restricted to lipid-based nanoparticles, and the larger shifts with HA-liposomes indicates that nanoparticle surface chemistry modulates the magnitude of *CTNNB1*-linked phenotype.

**Fig. 3.**
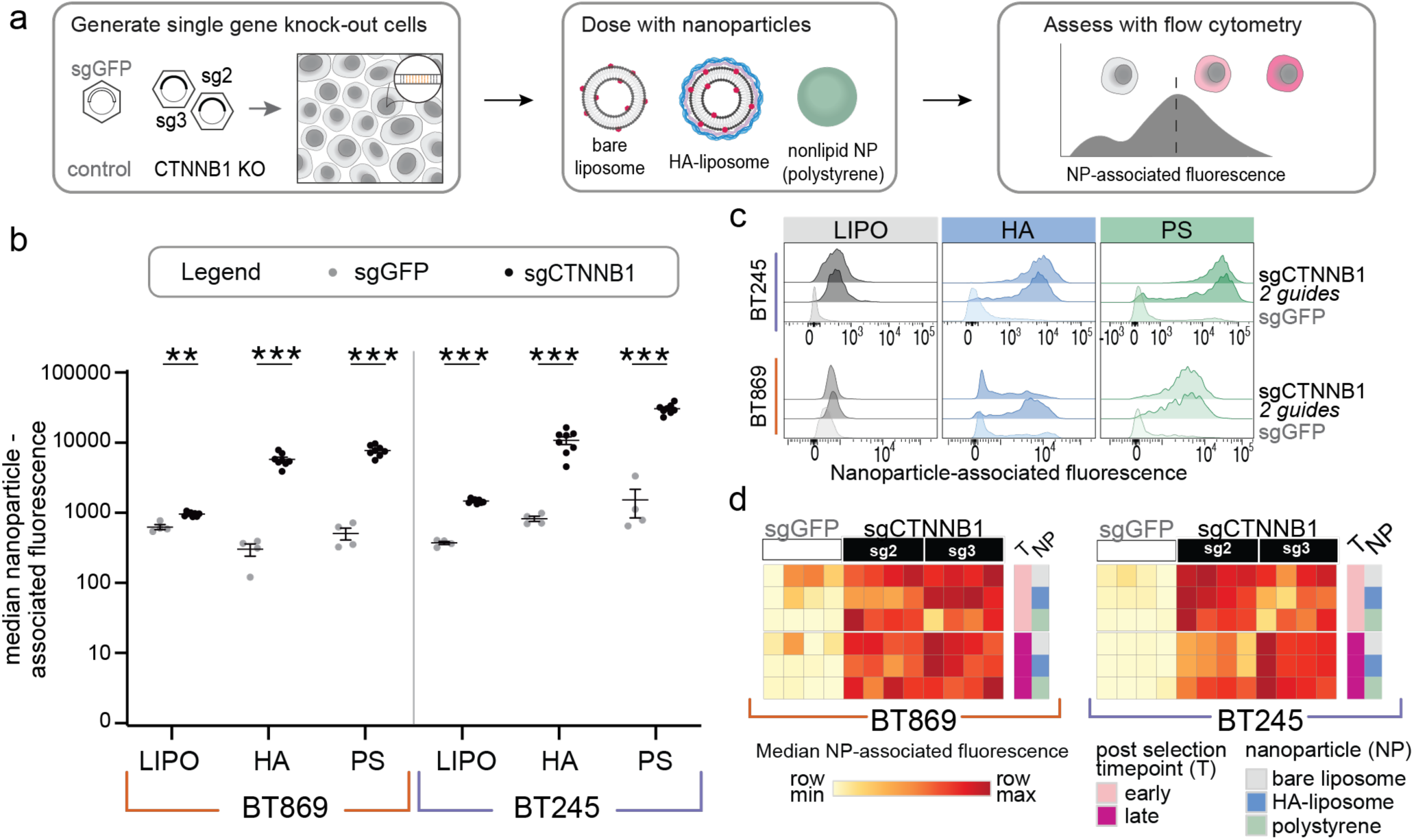
CTNNB1 knockout increases uptake of lipid and nonlipid nanoparticles in diffuse midline glioma models. **a**, Schematic of knockout model generation and sorting strategies following nanoparticle delivery. **b**, Raw values for the 4-hour incubation at late timepoint after antibiotic selection for both DMG models, data from one of two independent biological replicates. n = 4 (control) and n = 8 (knockout) technical replicates. The error bar represents mean ± standard error of mean; two-tailed Welch’s t-test was used for statistical analysis, ** indicates *p* < 0.01, *** indicates *p* < 0.001. **c**, Representative flow cytometry histograms from a late timepoint after antibiotic selection (day 11). **d**, Heatmap showing replicate- and guide-level nanoparticle association readout for 3 nanoparticle formulations assessed at 2 time points of nanoparticle incubation at early (1 day) and late (11 days) after antibiotic selection.

### CTNNB1 knockout changes cell membrane mechanics

Having observed that *CTNNB1* depletion increases both lipid and non-lipid nanoparticle uptake in DMG cells, we sought to define the molecular mechanisms underlying this observation. We reasoned that enhanced nanoparticle delivery following *CTNNB1* knockout could arise from two distinct but potentially overlapping roles of β-catenin including its canonical function as a transcriptional co-activator in Wnt signaling, or its structural role in cytoskeletal and membrane organization at the cadherin-based adherent junctions, and regulation of cellular uptake pathways (Fig. 4a). To investigate these possibilities, we performed RNA sequencing on *CTNNB1* knockout cells and control cells from the two abovementioned DMG models (BT245 and BT869), followed by Gene Set Enrichment Analysis (GSEA), using the C2:CP gene set database^33^ (Fig. 4b; Supplementary Fig. 7a-b). In the BT245 model, all the nine most-upregulated significant gene-sets across one or both guide RNAs were related to membrane-associated functions and structural remodeling in the setting of β-catenin loss. These included extracellular matrix (ECM) organization, cell-cell interactions, cytoskeletal maintenance, and transmembrane signaling, all of which exhibited positive normalized enrichment scores (NES) (Fig. 4b; Supplementary Fig. 7a). While not among the most significant depleted sets, we observed expected downregulation of canonical Wnt signaling following *CTNNB1* depletion, as evidenced by negative NES scores across majority of the pathways related to Wnt signaling (Fig. 4b; Supplementary Fig. 7a). These transcriptional changes suggest that in BT245 cells, *CTNNB1* depletion triggers not only the downregulation of canonical Wnt signaling, but also a shift toward membrane remodeling and structural reorganization.

Consistent with BT245, BT869 DMG cells also significantly downregulated canonical Wnt signaling following *CTNNB1* depletion (Supplementary Fig. 7b). We also observed dysregulation of three of the gene-sets related to ECM organization and interactions that were differentially expressed in BT245, including those related to extracellular matrix interactions, collagen formation and non-integrin membrane extracellular matrix interactions. However, while these pathways were observed to be upregulated in BT245, these specific pathways was significantly down-regulated following *CTNNB1* suppression in BT869 (Supplementary Fig. 7b), suggesting that *CTNNB1* loss may result in dysregulation of these processes through distinct mechanisms.

Based on these transcriptional changes, we hypothesized that *CTNNB1* loss would alter cellular mechanical properties. To test this, we measured single-cell stiffness using size-normalized acoustic scattering (SNACS), a microfluidic acoustic approach that reports on cell stiffness and is thought to be primarily sensitive to mechanical properties near the cell boundary^34^ (Fig. 4a and c, Supplementary Fig. 7). As expected, *CTNNB1* loss significantly reduced SNACS in both knockout populations at day 11 post-selection relative to controls in BT245 isogenic cell line model (Fig. 4c, *sgCTNNB1-2* vs *sgGFP*, p < 0.005; *sgCTNNB1-3* vs *sgGFP*, p < 0.0001). We observed similar trends in BT869, in which *CTNNB1* ablation and loss of β-catenin protein levels significantly reduced SNACS at day 7 (Supplementary Fig. 7d; *sgCTNNB1-2* vs *sgGFP*, < 0.0001; *sgCTNNB1-3* vs *sgGFP*, p < 0.0001). Western blot analysis for β-catenin performed concurrently confirmed efficient target suppression for both cell lines (Supplementary Fig. 7e-f). These data suggest that *CTNNB1* loss decreases cell stiffness across distinct DMG models, consistent with alterations in membrane and/or cortical mechanics.

Our results thus far suggest β-catenin ablation leads to membrane remodeling in DMG cells and prompted us to consider its role in endocytic activity. To further characterize the cellular phenotype associated with β-catenin loss, we leveraged 3D fluorescent microscopy to quantify receptor-mediated endocytosis and bulk phase macropinocytosis using transferrin and dextran as established macromolecular tracers, respectively, in *CTNNB1*-ablated cell models^35^ (Fig. 4d, Supplementary Fig. 8). Automated volumetric quantification of internalized puncta was performed, with integrated fluorescence intensity per cell representing a measure of endocytic or macropinocytic activity. Quantification of transferrin puncta confirmed a significant 1.6-fold (*sgCTNNB1-2*) and 1.9-fold (*sgCTNNB1-3*) increase in endocytic index in *CTNNB1*-ablated BT245 cells (*sgCTNNB1-2*, *p*<0.05 and *sgCTNNB1-3, p*<0.01, Fig. 4e). Conversely, quantification of dextran puncta revealed a significant 0.4- and 0.5-fold decrease in macropinocytic index in *CTNNB1*-ablated cells (*sgCTNNB1-2*, *p*<*0.001* and *sgCTNNB1-3, p<0.001,* Fig. 4f). This pattern is in line with prior evidence that the Wnt pathway positively regulates macropinocytic activity^36^, a process requiring dynamic structural re-organization to facilitate membrane ruffling and endosomal budding^37,38^, and suggests that its loss shifts the balance of endocytic activity in DMG cells.

Taken together, our transcriptomic, biophysical, and functional imaging data converge on a consistent picture: β-catenin loss in DMG cells results in reduced membrane stiffness, upregulation of membrane remodeling and ECM gene programs, and a shift in endocytic activity characterized by enhanced receptor-mediated endocytosis and reduced macropinocytosis. While the precise route by which nanoparticles exploit this altered membrane environment remains to be fully defined, the phenotypic profile is consistent with a cellular state more permissive to extracellular cargo uptake. These findings underscore the value of unbiased functional genomic screens in uncovering context-dependent biological regulators of nanoparticle delivery that would not be identified by hypothesis-driven approaches alone.

**Fig. 4.**
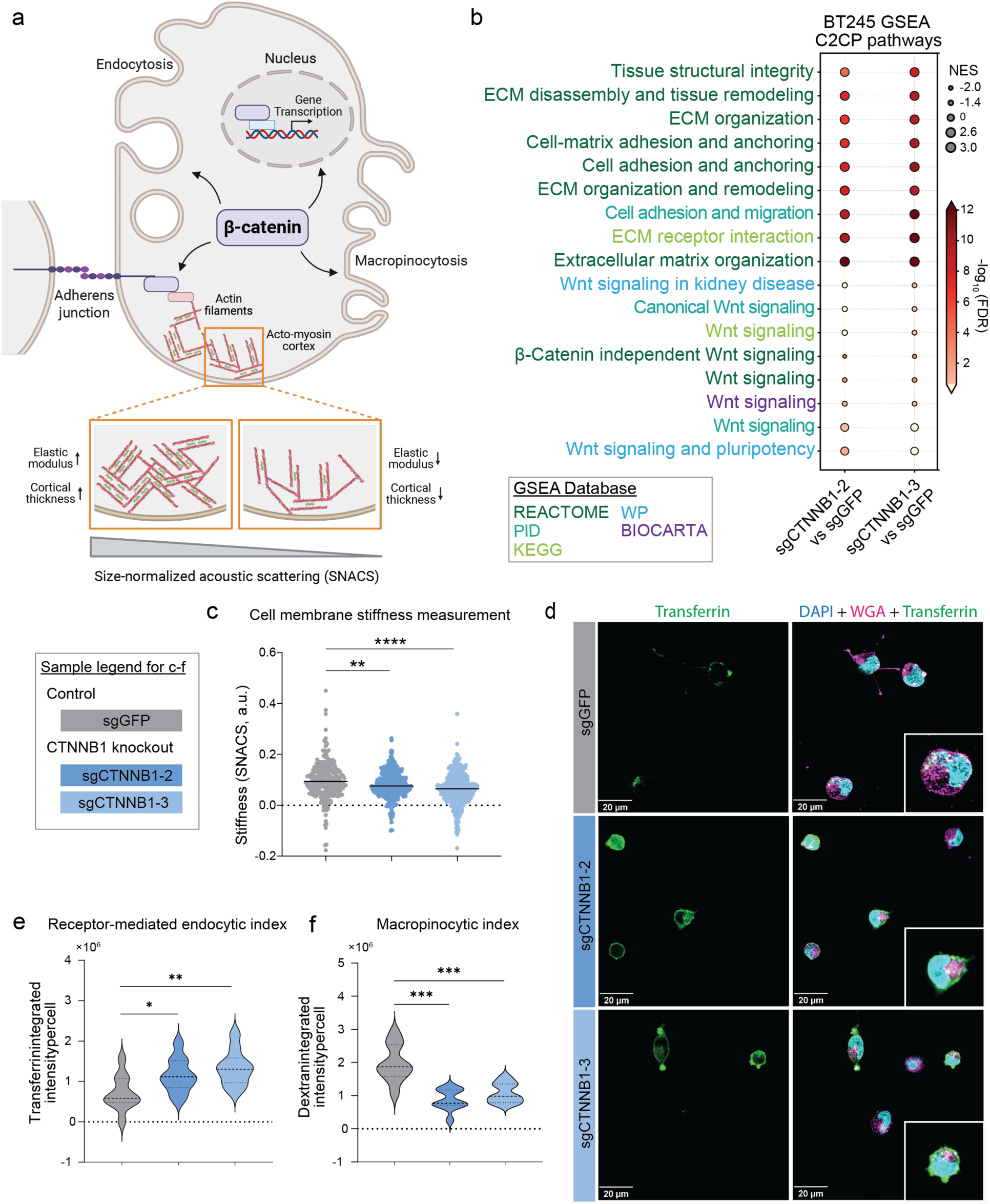
CTNNB1 knockout changes cell membrane mechanics and cellular uptake pathways in diffuse midline glioma models. **a**, Schematic figure showing canonical and non-canonical functions of β-catenin in the cell, emphasizing involvement of β-catenin in maintaining membrane structure and integrity. Schematic also illustrates how disruption of β-catenin leads to altered membrane mechanics that can be captured using size-normalized acoustic scattering (SNACS) representing membrane stiffness. **b**, Dot plot showing gene set enrichment analysis (GSEA) for CTNNB1 knockout cells using two independent guides in BT245 DMG model. The figure includes all significantly enriched and depleted C2:CP pathways below FDR 0.25., compared to control cell line. The pathways are ranked by NES. **c**, Violin plots showing size-normalized acoustic scattering (SNACS), a measure of single-cell stiffness, for control and CTNNB1 KO BT245 cell lines using two independent guides at day 11 post-selection. CTNNB1 depletion results in a significant reduction in cellular stiffness compared to control. Statistical significance was assessed using Welch’s t-test, with **** indicating *p* < 0.0001, and ** indicating *p* < 0.01. **d**, Z-projection of middle slices collected from 3D fluorescence microscopy of BT245 control and CTNNB1 knockout cells treated with transferrin, a receptor-mediated endocytosis probe. Violin plots showing normalized dot counting results of control and CTNNB1 depleted BT245 treated with (**e**) transferrin (for receptor mediated endocytosis) and (**f**) dextran (for micropinocytosis). Statistical significance was assessed using multiple Mann-Whitney test with Holm-Šídák correction, with * indicating *p* < 0.05, ** indicating *p* < 0.01, and *** indicating *p* < 0.001.

### MAPK pathway modulation of nanoparticle trafficking

Our pooled CRISPR-screen approaches also identified additional modulators of nanoparticle uptake beyond β-catenin, including multiple members of the MAPK and mTOR signaling pathways which were identified as negative regulators (Fig. 2b, Fig. 5a and b). Given that the MAPK axis is frequently upregulated across gliomas^39^, we reasoned that inhibition of this pathway may sensitize DMG cells to nanoparticle delivery. This was particularly of interest as several MAPK-pathway inhibitors have been approved for pediatric gliomas.

To test this combinatorial strategy, we pre-treated BT869 and BT245 cells with the MEK inhibitor trametinib followed by incubation with fluorescently labeled liposomal nanoparticles. Western blot analysis confirmed robust pathway suppression, as evidenced by a decrease in ERK phosphorylation (Supplementary Fig. 9a). MEK inhibition significantly increased nanoparticle association compared to DMSO-treated controls in both DMG models, although the magnitude was model-dependent. Following MEK inhibition, the BT869 cells exhibited a robust 3-fold increase (*p* < 0.01, Fig. 5c and d) in nanoparticle signal compared to vehicle control, whereas BT245 showed a modest 1.25-fold enhancement (*p* < 0.05, Supplementary Fig. 9b). To distinguish between surface attachment and true internalization of nanoparticles, we employed 3D fluorescence imaging. Upon observation of middle z-sections, with cell membrane labeled with wheat germ agglutinin, we confirmed that liposomes were located within the cytoplasmic compartment (Fig. 5e). This demonstrates that pharmacological inhibition of the MAPK pathway actively drives nanoparticle internalization in this context.

Collectively, these findings demonstrate the feasibility of deploying clinically relevant small molecule inhibitors to pharmacologically prime tumor cells for enhanced drug delivery. More broadly, this work provides a functional genomics framework for identifying, verifying, and optimizing context-dependent nanoparticle regulators.

**Fig. 5.**
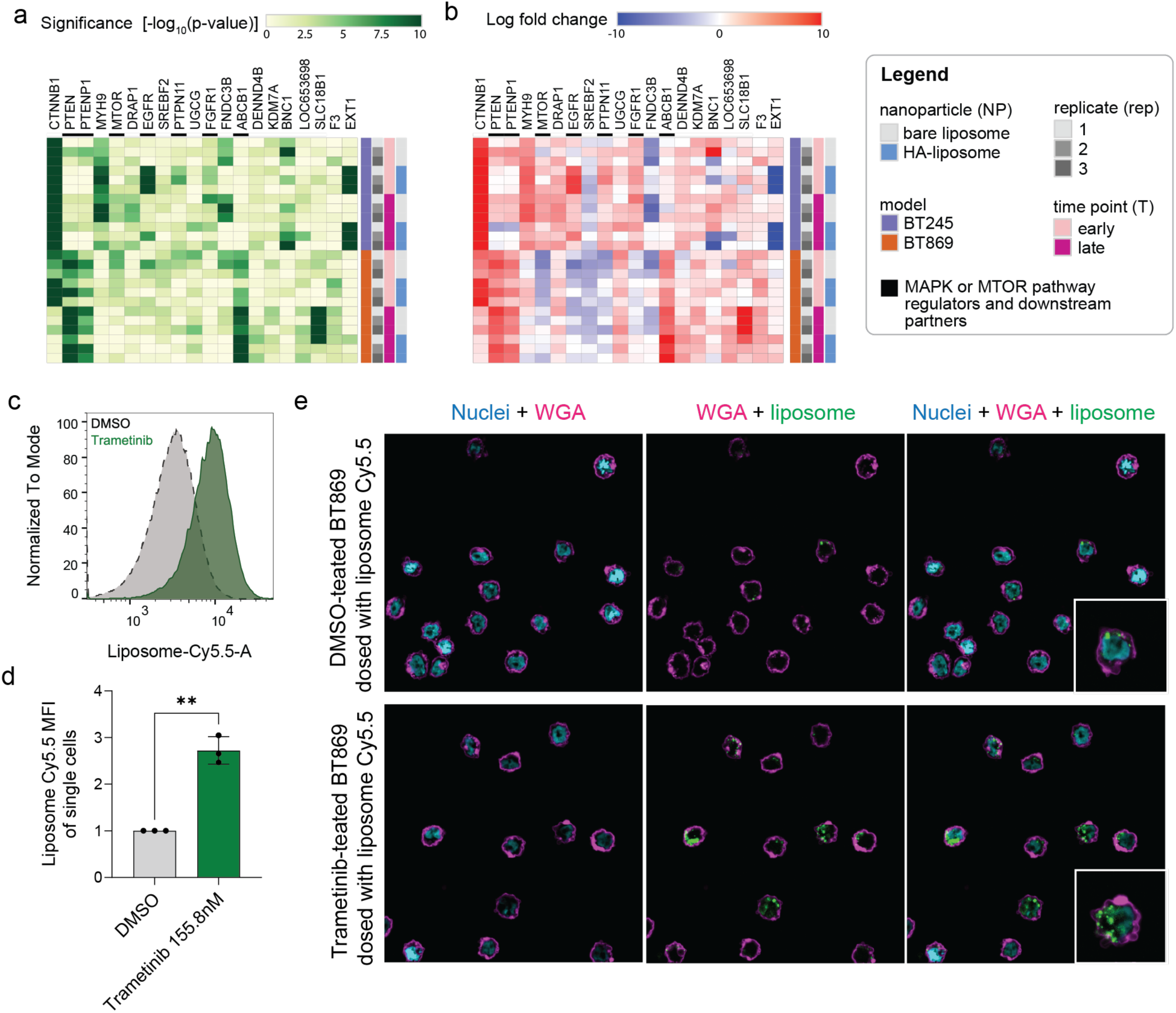
MAPK inhibitor modulates liposome uptake *in vitro*. Heatmap summarizing hits across screen conditions and top 20 candidate genes across all conditions by showing (**a**) statistical significance and (**b**) direction of expression changes, with positive log fold change (LFC) indicating guide barcodes being enriched in cells with high nanoparticle-association and negative LFC indicating guide barcodes being enriched in cells with low nanoparticle-association. (**c**) Representative histogram and (**d**) normalized MFI of liposome-association to MAPK inhibitor-treated BT869 measured with flow cytometry. (**e**) Fluorescence microscopy images on trametinib-treated BT869 after 24-hour of liposome dosing. Statistical significance was assessed using Welch’s t-test, with ** indicating *p* < 0.01.

## CONCLUSION

Genomic screens have transformed target discovery across cancer and other diseases, yet their application to nanomedicine remains limited. Designing and implementing a screen to effectively investigate the complex mechanisms governing physical interactions between a biomaterial and a cell requires a phenotypic readout as opposed to a fitness screen^40^. Here, we demonstrate that pooled genetic perturbation screens are a tractable and productive strategy to systematically uncover regulators of nanoparticle trafficking in clinically relevant pediatric brain tumor models.

The identification of β-catenin as a negative regulator of nanoparticle uptake was unexpected and mechanistically informative. Our data show that β-catenin levels influence membrane architecture and stiffness, with depletion leading to a cell state with enhanced extracellular cargo uptake. Whether this axis constitutes a predictive biomarker for nanomedicine responsiveness warrants additional investigation and may be especially meaningful if validated in the context of Wnt-driven tumors, as there are numerous therapeutics in development targeting the Wnt pathway.

Our finding that several members of the MAPK and mTOR signaling pathways modulate nanoparticle uptake is intriguing as there are several clinically relevant inhibitors for the treatment of pediatric gliomas. This raises the possibility that modulation of these signaling nodes could be combined with nanoparticle-encapsulated payloads for synergistic antitumor activity. This approach to combination therapy, in which one agent is employed to modulate the cell state and prime it for nanoparticle delivery, represents a new strategy for nanotherapeutics that may be amenable to rapid translation with existing clinical agents.

The screening framework described here is readily extensible along multiple axes. Expansion to genome-scale perturbation libraries, alternative tumor models, or endocytic phenotypes beyond surface association, such as endosomal escape, could further resolve regulatory architecture governing nano-bio interactions. More broadly, this approach is disease agnostic; any cellular system for which nanoparticle delivery can be quantified could be amenable to this methodology. While this work included two liposomal formulations, phenotypic screening across an extensive nanoparticle library across multiple cell types could provide the basis for rational nanoparticle design in the future. Ultimately, this platform shifts the development approach away from empirical, trial-and-error chemical modifications toward a predictive, biology-first screening methodology capable of systematically identifying, optimizing, and tailoring nanoparticle formulations.

## METHODS

### Cell lines

BT245 and BT869 DMG cell lines were obtained from the Center for Patient Derived Models at Dana-Farber Cancer Institute, courtesy of Dr. Keith Ligon. BT245 was cultured in tumor stem media containing Neurobasal-A (Gibco, 10888022) and DMEM/F-12 (Gibco, 11320082) at 1:1 ratio supplemented with 10mM HEPES buffer solution (Gibco, 15630080), 1x GlutaMAX supplement (Gibco, 35050061), 1nM sodium pyruvate (Gibco, 11360070), 1x MEM-NEAA (Gibco, 11140050), 1x Penicillin-Streptomycin (Gibco, 15140122), 0.5x B-27 supplement minus vitamin A (Gibco, 12587010), 0.0002% heparin (Stem Cell Technologies, 7980), 20 ng/mL EGF (Stem Cell Technologies, 78006.2) and 20 ng/mL bFGF (Stem Cell Technologies, 78003.2). BT869 was cultured in the same media with additional 7.5 ng/mL PDGF-AA (Fisher Scientific, 50990702) and 7.5 ng/mL PDGF-BB (Fisher Scientific, 50990710). Cells are seeded in ultra-low attachment flasks (Corning) at 100,000 – 200,000 cells per milliliter and passaged every 3-4 days.

### Antibodies and staining reagents

The following antibodies and dilutions were used for western blotting in this study: anti-β-catenin antibody 1:1,000 (Cell Signaling Technologies, 8480S and BD Biosciences, 610154), anti-β-actin-HRP antibody 1:8,000 (Santa Cruz Biotechnology, sc-47778), anti-phospho-p44/42 MAPK (Erk1/2) (Thr202/Tyr204) antibody 1:1,000 (Cell Signaling Technologies, 9101L), anti-p44/42 MAPK (Erk1/2) antibody 1:1,000 (Cell Signaling Technologies, 9102L), anti-vinculin-HRP antibody 1:10,000 (Santa Cruz Biotechnology, sc-73614), and anti-rabbit IgG-HRP antibody 1:5,000 (Cell Signaling Technologies, 7074). The following reagents were used for fluorescent microscopy sample staining in this study: mouse anti-β-catenin primary antibody 1:100 (BD biosciences,610154), donkey anti-mouse AF568 secondary antibody 1:200 (Thermo Fisher, A10037), wheat germ agglutinin-AF633 (Thermo Fisher, W21404), wheat germ agglutinin-AF488 (Thermo Fisher, W11261), DAPI (Thermo Fisher, D1306), and Hoechst 33342 (Thermo Fisher, H1399).

### Pooled viral library production

Virus for the pooled library was generated by the Broad Institute Genetic Perturbation Platform. Virus for single gene knock-out experiments was produced in house. Briefly, HEK293T cells were plated in 10 cm dishes (Celltreat Scientific Products). Once cells reached 90% confluence, they were transfected with 10 µg of psPAX2, 1 µg of VSVG, and 10 µg of lentiviral plasmid of interest, utilizing Lipofectamine 3000 (Thermo Fisher, L3000015). HEK293T cell media was changed after six hours. 24 hours following transfection, the supernatant containing lentivirus was collected with Luer-Lok syringes (Fisher Scientific, 302995) and filtered utilizing 0.45 µm PES syringe filters (Fisher Scientific, SLHPR33RS). Lentivirus was concentrated using Lenti-X Concentrator (Takara, 631232) according to the manufacturer’s instructions.

### Screen design

An all-in-one CRISPR knockout vector was chosen with Cas9 and guide RNA on the same plasmid backbone (pXPR_023). 705 genes of interest were selected based on previously identified candidate biomarkers^14^ and genes related to cell membrane binding and endolysosomal trafficking. Four guides were designed for each gene, and an additional 150 intergenic and 150 non-targeting controls were also included. A total of 22 genes included in the screen (3%) are categorized essential genes. To control for background effects, the library also included 300 non-targeting or intergenic region targeting sgRNAs. The screen was designed in collaboration with the Broad Genetic Perturbation Platform (GPP). In Fig. 2, each dot represents the average log-fold change of across four guides targeting each gene and include three independent replicates. Volcano plots for each line are included at early and late timepoints for each nanoparticle (bare liposome and HA-liposome).

### Infection and selection of the pooled lentiviral library

DMG cell lines were infected in 12-well plates (Corning) at a density of 3 million cells per well. Each transduction was done in triplicate with one well per replicate. No-infection controls were included on the same plate for all infections. DMG cell lines were infected by spinfection at 2000 rpm and 30°C for two hours. 18 hours after spinfection, cells were detached from the plates using Accutase Enzyme Cell Detachment Medium (Thermo Fisher, 501129055), seeded into ULA plates (Corning), and subjected to puromycin selection (0.75 µg/mL for BT245, 0.5 µg/mL for BT869) for 48 hours before downstream assays.

### Genomic DNA preparation, sequencing, and guide enrichment analysis

For FACS based CRISPR screens, cell pellets were collected for each sorted population and maintained at -80°C until genomic DNA preparation. NucleoSpin Blood Mini kits (Takara, 740951.5) were used according to manufacturer instructions with one modification. After cell lysis, DNA precipitation, and DNA binding steps, the elution step was performed overnight at 4°C before centrifuging. PCR inhibitor removal was then performed using OneStep PCR Inhibitor Removal Kit (Zymo, Z-D6030) according to manufacturer instructions. DNA concentration was measured using the Qubit dsDNA HS Assay Kit (Invitrogen, Q32851) according to manufacturer instructions. PCR was used to amplify the LentiCRISPRv2 barcodes. Samples were submitted for targeted sequencing of barcodes corresponding to each guide, alongside with a plasmid DNA control. Apron (Broad Institute GPP) was used to analyze the distribution of each guide relative to the plasmid DNA, enabling enrichment/depletion measurements.

### Fluorescent activated cell sorting (FACS)

For phenotypic CRISPR screens, cells were incubated with fluorescent nanoparticles at a final well concentration of 0.01 mg/mL for a set time period (1, 4, or 24 hours). Cells were then washed once with PBS and treated with Accutase Enzyme Cell Detachment Medium (Thermo Fisher, 501129055) for 5 minutes to obtain a single cell suspension and transferred to a FACS tube through a cell strainer cap (Falcon). Samples were stored on ice until analysis, and processed within 2 hours of preparation. Using a BD FACSAria II Cell Sorter (BD Biosciences), single cells populations were identified and sorted based on APC signal (excitation 640, filters 660/20). We use a dynamic gating strategy to collect the lowest and highest quartile for each sample.

### Nanoparticle formulation and characterization

Lipids were obtained from Avanti and dissolved in chloroform, methanol and water in a 65:35:8 volume ratio mixture. Non-PEGylated, fluorescent liposomes were generated with 1,2-distearoyl-sn-glycero-3-phosphocholine (DSPC, 850365C), 1,2-distearoyl-snglycero-3-phospho-(1’-rac-glycerol) (DSPG, 840465), cholesterol (700100), 1,2-distearoyl-sn-glycero-3-phosphoethanolamine (DSPE, 850715), and DSPE-Cy5 (810345C) or DSPE-Cy5.5 (810336) at 31:31:6.8:0.2 molar ratio as previously described^14^. Briefly, the lipid solution was evaporated using a rotovap system until completely dry, then rehydrated at 60°C under sonication with milliQ water. Liposomes were extruded through sequentially smaller membranes until a uniform population between 80-100 nm in diameter was formed. Layer-by-layer liposomes were generated by adsorbing positively charged poly-L-arginine hydrochloride (PLR200, 38.5 kDa, Alamanda Polymers), purifying excess polymer through tangential flow filtration, and adsorbing sodium hyaluronate (HA40K-1, HA, 40 kDa, Lifecore Biomedical) using established prior methodology^14,41^ and weight ratios (polymer to lipid) of 0.8:1 and 1:1, respectively. Polystyrene nanoparticles were sourced commercially as 100 nm carboxylated FluoSpheres (Invitrogen), in the yellow-green color. All nanoparticles were characterized using dynamic light scattering (Malvern ZS90 Particle Analyzer) for hydrodynamic size and polydispersity. Using the same instrument and laser doppler electrophoresis, surface zeta potential was monitored before and after each layering step and prior to downstream experiments.

### β-catenin knockout cell line preparation

HEK293T cells were transfected with *sgCTNNB1-2*, *sgCTNNB1-3*, or *sgGFP* control transfer plasmids, psPAX2 packaging plasmid, and pMD2.G envelope plasmid using Lipofectamine P3000 (Thermo Fisher, L3000015) in OptiMEM (Fisher Scientific, A4124801) according to manufacturer instructions. Lentivirus was concentrated from 10 mL to 1 mL using Lenti X concentrator (Takara Bio, 631232) according to manufacturer instructions.

Puromycin (Thermo Fisher, A1113803) titration was performed on BT245 cells. 1.25 μg/mL puromycin was chosen by reducing the wild type cell viability to 0%. Viral titration was performed on BT245 cells by adding a range of concentrations for each guide, and following the steps outlined below. 3 million BT245 cells were seeded per guide with one additional non-infection control (NIC) well in a tissue-culture treated 12-well plate. Media and virus or PBS for NIC were added to a total volume of 2 mL. Cells were then centrifuged for 2 hours at 2000 rpm on a swing bucket centrifuge. Immediately after spin, 2 mL media was added to each well. The plates were incubated for 24 hours at 37°C and 5% CO2. Cells were collected and treated with 100 μL Accutase (Thermo Fisher, 501129055) to achieve single cell suspension. Cells were gently pipette-mixed with a 200 μL pipette tip for 3 minutes, diluting with 900 μL media. 1 mL infected cells were then transferred to 3 mL media per well in a ULA 6 well plate (Corning, 3471). 1 mL of puromycin (Thermo Fisher) with 5 times the selected concentration (i.e., 1.25 μg/mL) was added to each well. Cells were incubated in puromycin media for 72 hours before counting and confirming 0% viability in the NIC well. Transduced cells were returned to culture at 200,000 cells/mL until their viability returned to 90%.

### Cloning CRISPR-Cas9 LVV transfer plasmids

Circular all-in-one CRIPSR KO vector (lentiCRISPRv2; pXPR_023), which contains a Cas9 nuclease and customizable sgRNA expression cassette, was obtained from the Broad Institute Genetic Perturbation Platform (GPP). The circular vector was linearized using BsmBI (New England Biolabs, R0739L) according to manufacturer instructions. The guide sequences targeting GFP (control) and β-catenin were then cloned into the plasmid DNA backbone, selected/amplified with *E. coli* growth, and purified for LVV production. Guide sequences are listed as follow: TTTACGTCGCCGTCCAGCTC (*sgGFP* sequence), CTGGGACCTTGCATAACCTT (*sgCTNNB1-2* sequence), and GCTTATTACTAGAGCAGACA (*sgCTNNB1-3* sequence).

### Western blotting

1x RIPA/protease inhibitors were made from 10x RIPA Buffer (Cell Signaling Technology, 98065) and 100x Halt Protease and Phosphatase Inhibitor Cocktail (Thermo Fisher, 78440). Cell pellets were lysed with 1X RIPA/PI while rotating at 4°C for one hour. Lysates were centrifuged at 17,000 x g for 15 minutes at 4°C and the supernatant was collected. Protein quantification was performed using Pierce 660 nm Protein Assay Reagent (Thermo Fisher, 22660) with samples diluted 1:5. Five technical replicates of diluted supernatant were added to a 96-well plate (Fisher Scientific, 12565501) for each sample. Plates were read on a Molecular Devices SpectraMax M5 plate reader.

Lysates were mixed with 10X NuPAGE Sample Reducing Agent (Thermo Fisher, NP0004) and 4x NuPAGE LDS Sample Buffer (Thermo Fisher, NP0007) before heating at 95°C for 5 minutes. The samples were loaded and run on NuPAGE 4 to 12% Bis-Tris 1.5 mm Mini Protein Gels (Thermo Fisher, NP0335) at a constant voltage of 140 V for 90 minutes. Amersham ECL Full-Range Rainbow Molecular Weight Marker (Cytiva, RPN800E) was used a ladder on each gel. The iBlot 2 Dry Blotting System (Thermo Fisher, IB21001) was used for dry transfer of the blots onto PVDF membranes (Thermo Fisher, IB24001). In all subsequent steps, 5% milk in TBST was utilized for blocking and antibody dilution. Following transfer, blots were blocked at room temperature for one hour and then incubated in primary antibody overnight at 4°C. Blots were washed three times in TBST (10 minutes each), followed by incubation in secondary antibody at room temperature for one hour and another wash with TBST. SuperSignal West Femto Maximum Sensitivity Substrate (Thermo Fisher, 34095) was used for chemiluminescent imaging with a GE ImageQuant LAS 4000 imager.

### Cell viability assay

Cells were seeded at a density of 1,000 cells per well in a 96-well plate (Corning, 3917), and cell proliferation was measured for 6 consecutive days. The CellTiter-Glo 2.0 viability assay (Promega, G7573) was used to measure the level of ATP as a surrogate for cell viability. Following addition of CellTiter-Glo reagent, plates were incubated and gently shaken for 10 minutes in the dark to ensure effective cell lysis. Luminescence was measured in a SpectraMax M5 plate reader (Associated Technologies Group) with the SoftMax Pro software using the CellTiter-Glo protocol with an integration time of 500 ms.

### Incucyte spheroid assay

The Incucyte Live-Cell Analysis System (Sartorius) was utilized for *in vitro* cell proliferation assays. DMG cell lines were plated in ultra-low attachment (ULA) 96-well round-bottom plates (Corning) at a density of 1,000 cells per well. Plates were spun at 200 x g for 15 minutes at room temperature to initiate spheroid formation at the bottom of the wells. Time-lapse imaging of spheroid size was obtained every six hours for seven days using the spheroid module. Spheroid sizes were measured using the Incucyte software by analyzing the largest brightfield object.

### RNA sequencing and analysis

Bulk RNA was extracted from BT245 and BT869 isogenic cell lines (8 samples for BT245 isogenic sets, 9 samples for BT869 for BT245 isogenic sets, 2-3 technical replicates were used for each sample) using the RNeasy mini kit (Qiagen, 74104). 1 µg RNA for each sample was submitted to the MIT BioMicro Center. RNA was subjected to sequencing using AVITI24 from Element Biosciences, leveraging the NEB Ultra II kit for library preparation, and sequencing to produce 75 base pair paired-end reads.

Raw RNA-sequencing reads were processed using the nf-core/rnaseq pipeline (version 3.14.0). Reads were aligned to the hg38 human reference genome and quantified against the GENCODE (v43) basic gene annotation using the STAR-Salmon (star_salmon) alignment and quantification mode. Transcript-level abundances were summarized to gene-level counts using the tximport R package (v1.32.0). Raw and normalized counts matrices, expressed as Counts Per Million (CPM), were constructed independently for each cell line. Genes were filtered from the count matrices if they failed to reach an expression level of at least 0.3 CPM in three or more samples. Following this filtering step, 17,108 and 18,115 genes were retained across the BT245 and BT869 samples, respectively. Differentially expressed genes (DEGs) were identified using the DESeq2 R package (v1.44.0). For each cell line, sgCTNNB1 knockout replicates (sgCTNNB1-2 and sgCTNNB1-3) were compared against the sgGFP control. The output from DESeq2 was first filtered to retain only protein-coding genes based on GENCODE v43 annotations. Genes were then ranked in descending order by their Wald statistic to generate a ranked gene list, which was used for gene set enrichment analysis (GSEA) with the fgsea R package (v1.30.0). Human gene sets were retrieved from the Molecular Signatures Database (MSigDB) via the msigdbr R package (v26.1.0), including the C2:CP (Canonical Pathways) collections. Gene sets containing fewer than 15 or more than 500 genes were excluded from the analysis. GSEA results were first filtered to retain significantly dysregulated pathways (FDR < 0.25) in at least one of the two CTNNB1-targeting guide-level comparisons. This list was further narrowed to pathways related to WNT signaling and ECM/membrane remodeling, and the shortlisted pathways were visualized in a dot plot generated with matplotlib.pyplot (v3.10.1).

### Fluorescent 3D microscopy sample preparation and imaging

LabTek II borosilicate 8-well chamber slides (Thermo Fisher, 155409) were coated with 50 μg/mL rat tail collagen (Corning, 354236) in 0.02 N acetic acid. For macromolecule uptake mechanism studies, BT245 with and without *CTNNB1* knockout were seeded at 70,000 cells per well in media. Either 0.01 mg/mL 70,000 MW lysine-fixable tetramethylrhodamine (TMR)-dextran (Thermo Fisher) or 0.04 mg/mL transferrin in complete media was then added to the wells and incubated for 30 minutes. The cells were washed once with cold PBS before being fixed with 4% formaldehyde. The cells were then permeabilized for 15 minutes with 300 μL 0.1% Triton X (Thermo Fisher) and blocked overnight in 5% milk in TBS-T at 4°C. The next day, cells were washed and stained with 2.5 μg/mL mouse anti-β-catenin primary antibody (BD biosciences) for one hour at 4°C, followed by another PBS wash before staining of 10 μg/mL donkey anti-mouse AF568 secondary antibody (Thermo Fisher) for one hour at 4°C. To visualize cell membrane and nuclei, the cells were sequentially stained with 5 μg/mL wheat germ agglutinin-AF633 (Thermo Fisher) and 2 μg/mL DAPI (Thermo Fisher).

For evaluating internalization of liposomes, BT869 cells were seeded in ultra-low attachment 12 well plate at 1.5 million cells per milliliter and treated with either 155.8 nM trametinib (MedChemExpress, HY-10999) or 0.01% DMSO for 24 hours. Next day, the cells were washed, treated with Accutase, and re-seeded into a new plate at 150,000 cells per well for liposome dosing. 20 µg/mL lipid blend of Cy5.5-liposome were added to the culture for another 24-hour incubation. On the day of sample staining, all groups were treated with Accutase, washed with PBS, and fixed with 4% formaldehyde. To visualize cell membrane and nuclei, the cells were sequentially stained with 5 μg/mL wheat germ agglutinin-AF488 (Thermo Fisher) and 5 μg/mL Hoechst 33342 (Thermo Fisher) in Eppendorf tubes. The cells were then resuspended in PBS and let settle in a LabTek II chamber slide well before slow aspiration of PBS and mounting.

ProLong Diamond (Thermo Fisher, P36961) were used to mount all samples. Cells were imaged using a Leica DMi8 microscope with Thunder deconvolution software (LAS X, Leica Microsystems) and a 100x oil immersion objective. Z-stack images were taken with 0.5 μm step-size and computationally cleared using either the Small Volume Computational Clearing (uptake mechanism) or Instant Computational Clearing (liposome internalization), and Prolong Diamond refractive index settings.

### Cell morphology grading

For initial preprocessing of the images collected using the Leica DMi8 microscope and the Thunder system, we used ImageJ for converting LIF files to TIFF format and splitting individual channels. The TIFF files were then processed with CellProfiler and ImageJ software for cell count and speckle count quantification, respectively. The bottom 4 slices were removed to avoid collecting background signal from the coated glass. DAPI and WGA fluorescence signal, processed with CellProfiler rescaling, median filtering, closing, and watershed functions, were used to identify and separate individual cells. After size filtering on ImageJ, z-projected standard deviation of fluorescently labeled transferrin and dextran were used to locate endosome and macropinosome speckles, respectively. Integrated fluorescent signal across z-slices (from slice 5) was used for uptake index calculation. Total speckle fluorescence intensity of each 3D image stack was then normalized by cell number.

### Flow cytometry particle association assay

DMG cell lines were treated with Accutase for 4-7 minutes. They were then seeded into an ultra-low attachment 6 well plate at 1-2 million cells per milliliter and treated with trametinib or DMSO for 24 hours. The treated cells were harvested and seeded into a new ultra-low attachment well plate at 150,000 cells per milliliter for incubation with liposomes. The cells were either dosed with liposome-Cy5.5 for 4 hours or 24 hours at 10 µg lipid blend per milliliter. To prepare cells for flow cytometry, they were spun down and treated with Accutase again, followed by an additional wash with PBS with 1% BSA (Miltenyi, 130-091-376) for removing non-specific association of liposomes. The cells were then filtered with 30 µm filters to avoid clumping. The cells were kept at 4°C prior to sample loading on NovoCyte cytometer (Agilent).

### Acoustic scattering measurements

Size-normalized acoustic scattering (SNACS) was carried out using an established microfluidic methodology^34^. In brief, cells in single cell suspension were flowed through an acoustic wave generated by a vibrating suspended microchannel resonator (SMR)^42^. In parallel, cell volume was measured using a Coulter Counter (Beckman Coulter). The average cell volume was determined and used to calculate SNACS on a per cell basis. Before each experiment, the SMR was cleaned before use with 0.25% Trypsin-EDTA for 20 min, then 5% bleach for 2 min, then DI-H_2_O for 2 min to remove any biological debris. After cleaning, a solution of 1 mg/mL PLL-g-PEG was applied at room temperature for 10 min to passivate the SMR. Finally, it was rinsed with cell culture media before and after each measurement. Latrunculin B (lat-B) was used to treat control (nontransduced) cells as a positive control to decrease cell stiffness. Sample loading was through 0.005-inch-inner-diameter fluorinated ethylene propylene (FEP) tubingand three independent electronic pressure regulators (MPV1, Proportion Air) were used to drive fluid flow. Pressure regulation and data acquisition were controlled using custom software written in LabVIEW (National Instruments). Data analysis was performed in MATLAB R2022b.

## ACKNOWLEDGEMENTS

We thank the CPDM team and Dr. Keith Ligon for providing the DMG cell lines used in this study. We also thank the Koch Institute’s Robert A. Swanson Biotechnology Center for technical support, specifically the flow cytometry core. Figure 4a is created using BioRender and publication license has been obtained (Beroukhim, R. (2026) https://BioRender.com/ckxaaq9).

## AUTHOR CONTRIBUTIONS

Investigation: E.L.C., S.P., J.W.T., C.B.Y., J.K-S., M.C., M.O., K.Y.C., E.V., E.B., K.R.B., M.M.B., C.A.W., Y.Z., M.D., C.F.

Conceptualization: P.B., J.P.S

Methodology: S.B., Y.Z., M.D., N.K., J.G.D.

Formal analysis: S.R., C.A.W., Y.B., J.P.S.

Writing – original draft: E.L.C., S.P., J.W.T., P.B., J.P.S.

Writing – review & editing: E.L.C., S.P., P.B., J.P.S.

Supervision: K.L.L., S.R.M., P.T.H., P.B., J.P.S.

Funding acquisition: P.T.H., P.B., J.P.S.

