## Supplementary Information for "Phenotypic screens identify biologic regulators of nanoparticle uptake in diffuse midline glioma"

**Supplementary Fig. 1** Flow cytometry gating strategy and technical repeat data of nanoparticle CRISPR-Cas9 screen cell sorting.

**Supplementary Fig. 2** Quality control of CRISPR-Ca9 knockout library guide representation.

**Supplementary Fig. 3** Number of candidate genes identified through CRISPR-Cas9 screen, categorized based on –log(p), in DMG models with different nanoparticle incubation durations.

**Supplementary Fig. 4** Concordance and discord of CRISPR-Cas9 screen hits between the DMG models.

**Supplementary Fig. 5** *CTNNB1* single guide RNA selection and gene dependency in DMG cell lines.

**Supplementary Fig. 6** Raw median fluorescence intensity data for *CTNNB1* knockout experiments in BT245 and BT869 models for incubation times not shown in Fig. 3D.

**Supplementary Fig. 7** *CTNNB1* knockout changes cell membrane mechanics in diffuse midline glioma models.

**Supplementary Fig. 8** Z-projection of middle slices collected from 3D fluorescence microscopy of BT245 control and *CTNNB1* knockout cells treated with dextran, a macropinocytosis probe.

**Supplementary Fig. 9** MEK inhibitor validation in both DMG models and nanoparticle association in BT245.

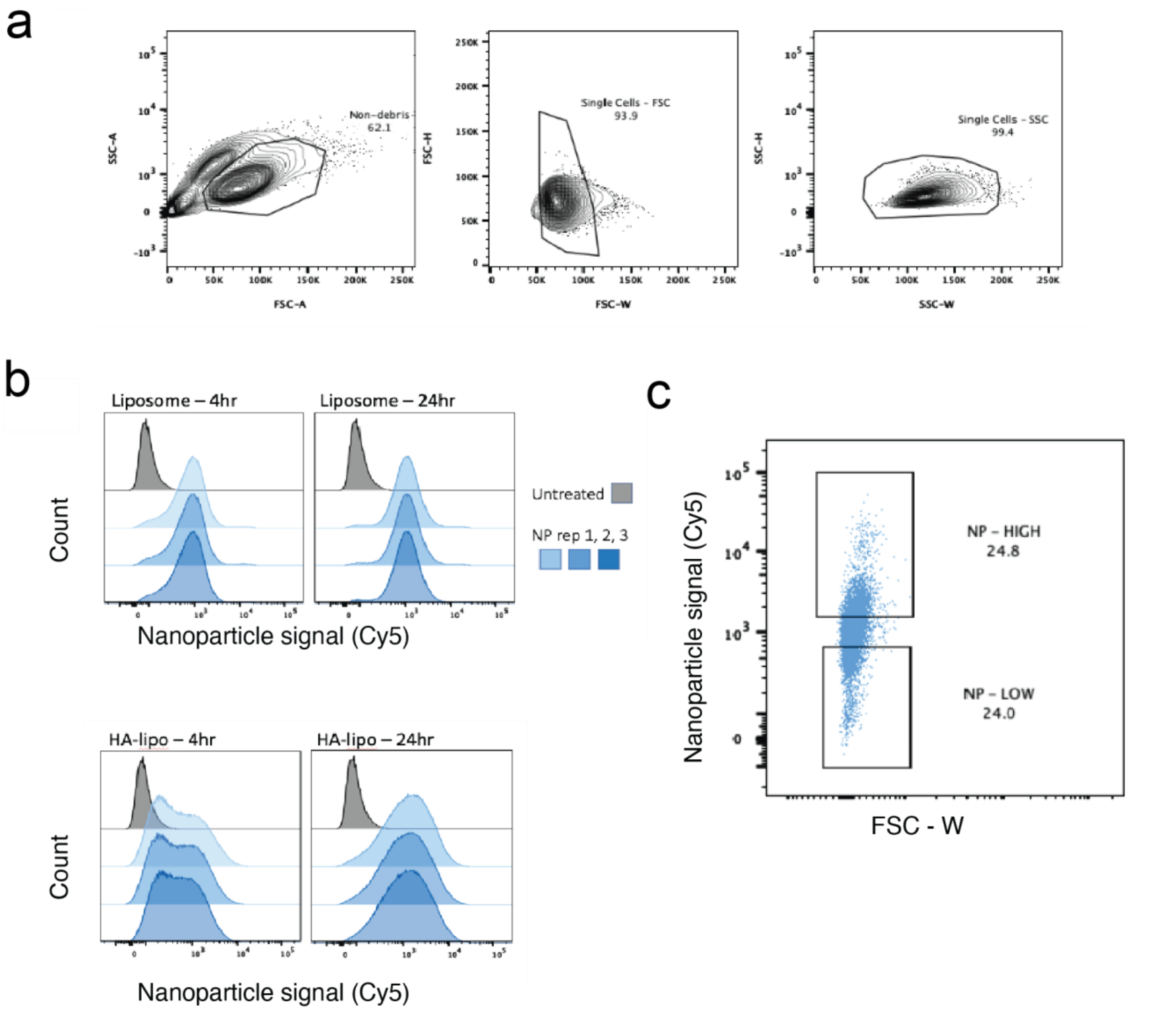


**Supplementary Fig. 1 | Flow cytometry gating strategy and technical repeat data of nanoparticle CRISPR-Cas9 screen cell sorting. a**, Gate setting for non-debris single cells. **b**, 3 technical replicate histograms of bare- and HA-liposome fluorescent signal after 4- and 24-hour incubation with transduced DGM cells. **c**, Gating the top and bottom quartile of nanoparticle-associated knockout library transduced-DMG populations.

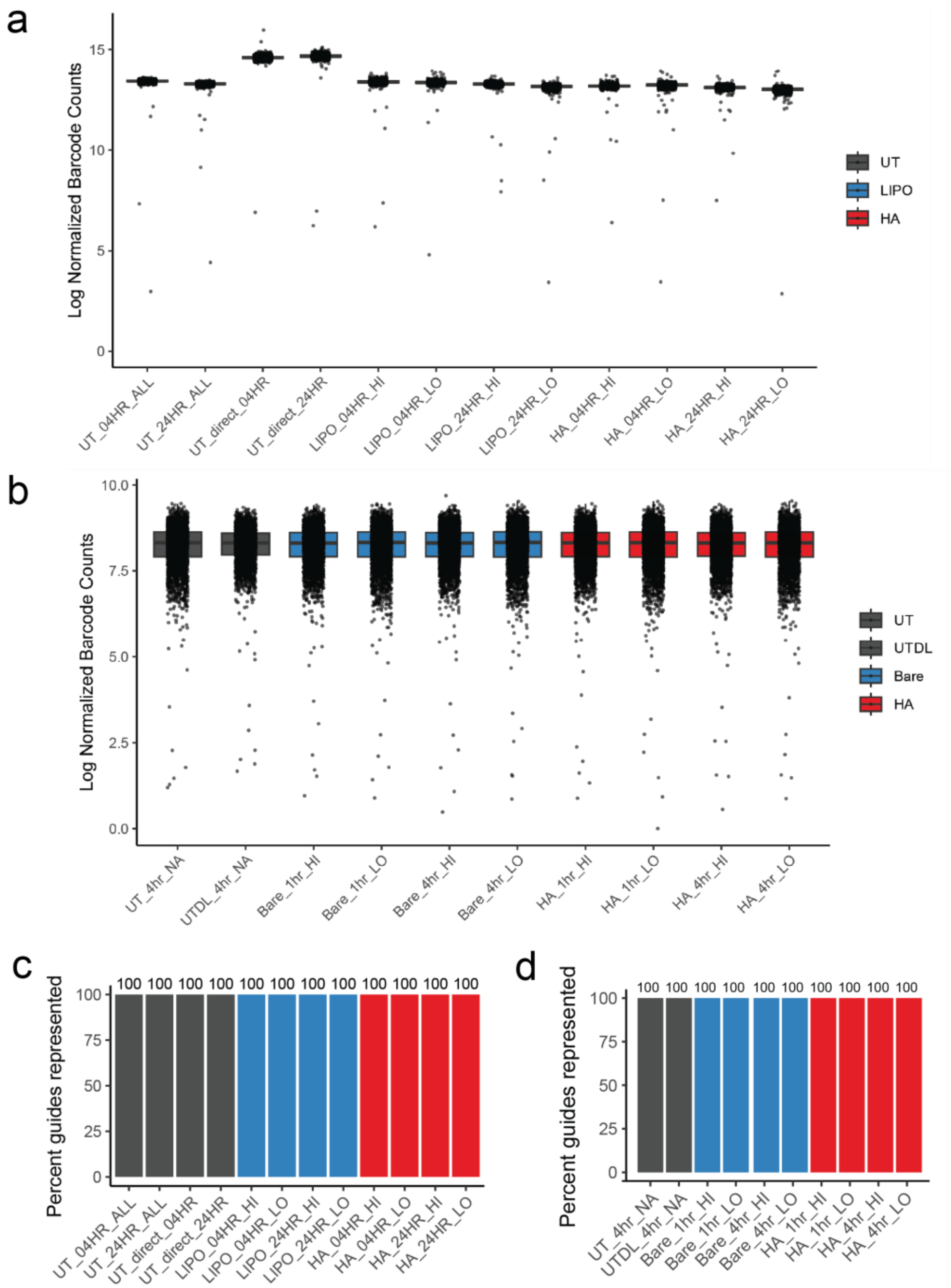


**Supplementary Fig. 2 | Quality control of CRISPR-Ca9 knockout library guide representation.** Knockout library barcode count in (**a**) BT245 and (**b**) BT869, as well as percentage guide representation in (**c**) BT245 and (**d**) BT869 populations with different degree of nanoparticle association after 1-, 4-, or 24-hour incubation.

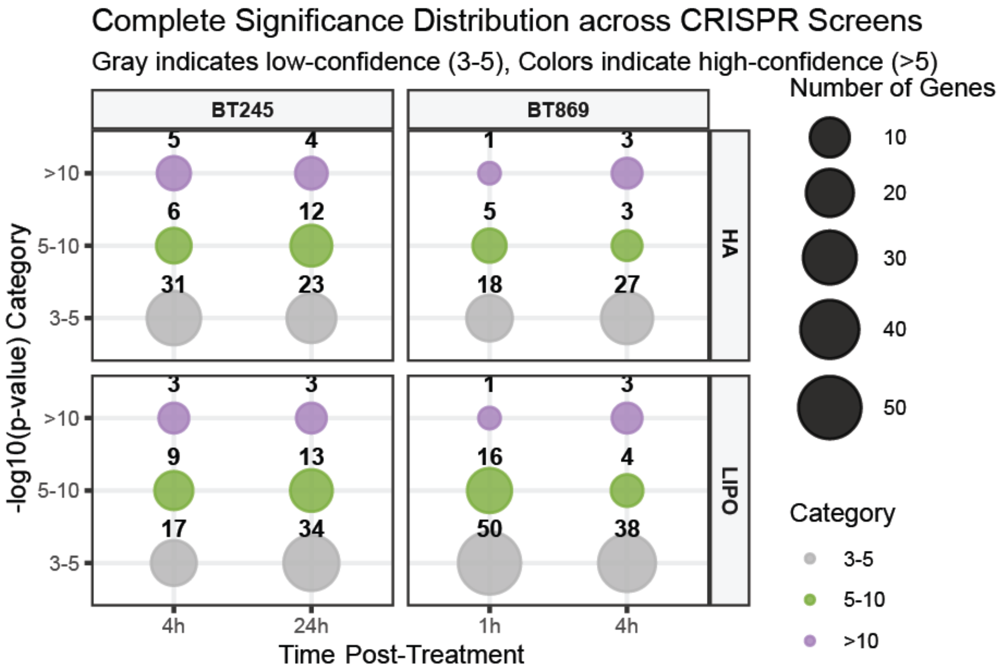


**Supplementary Fig. 3 | Number of candidate genes identified through CRISPR-Cas9 screen, categorized based on –log(p), in DMG models with different nanoparticle incubation durations.**

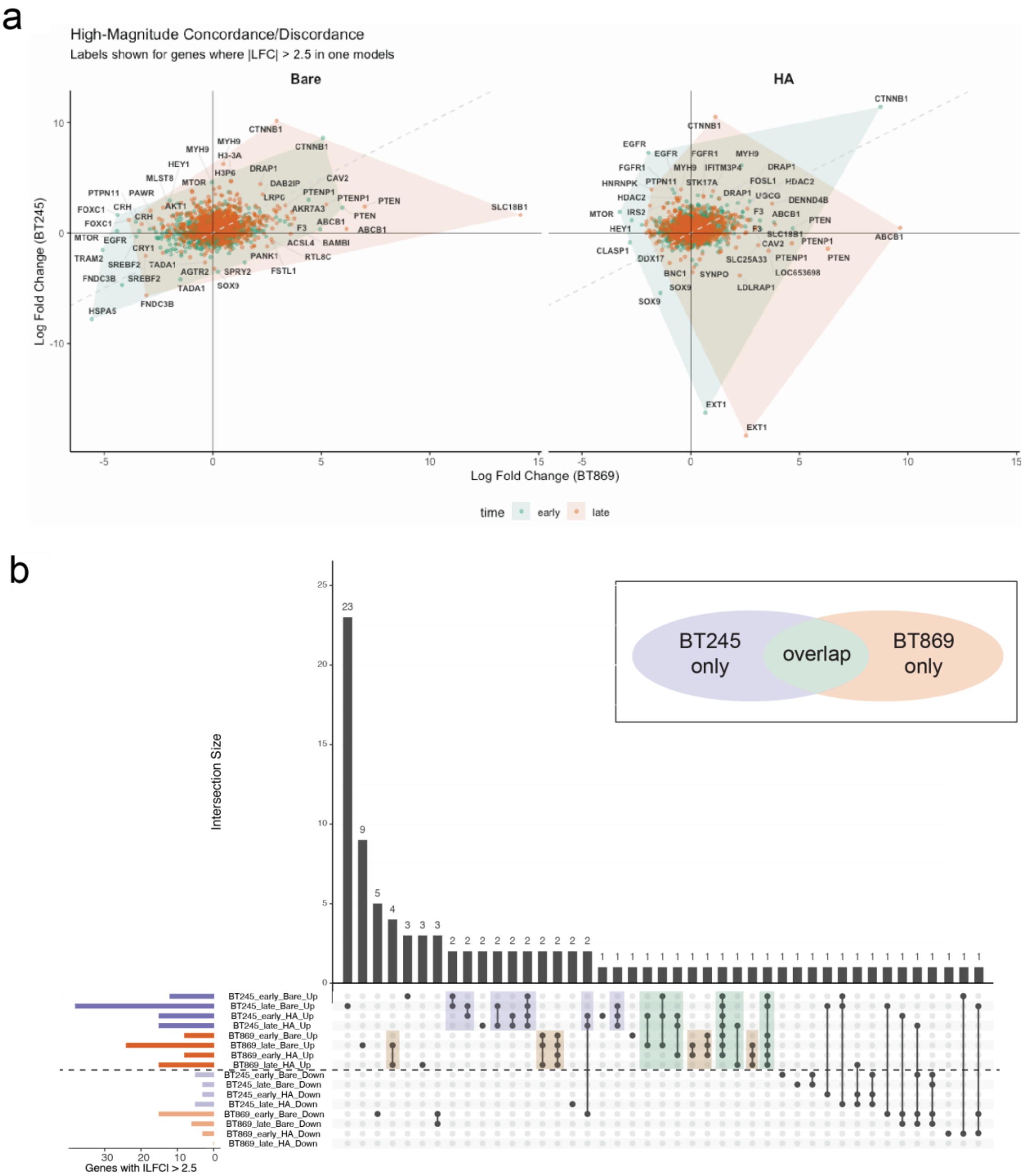


**Supplementary Fig. 4 | Concordance and discordance of CRISPR-Cas9 screen hits between the DMG models. a**, Overlap and diverge hits with |LFC| > 2.5 in at least one of the DMG models. **b**, Upset plot of the top CRISPR hits with change direction in BT245 and BT869 cells.

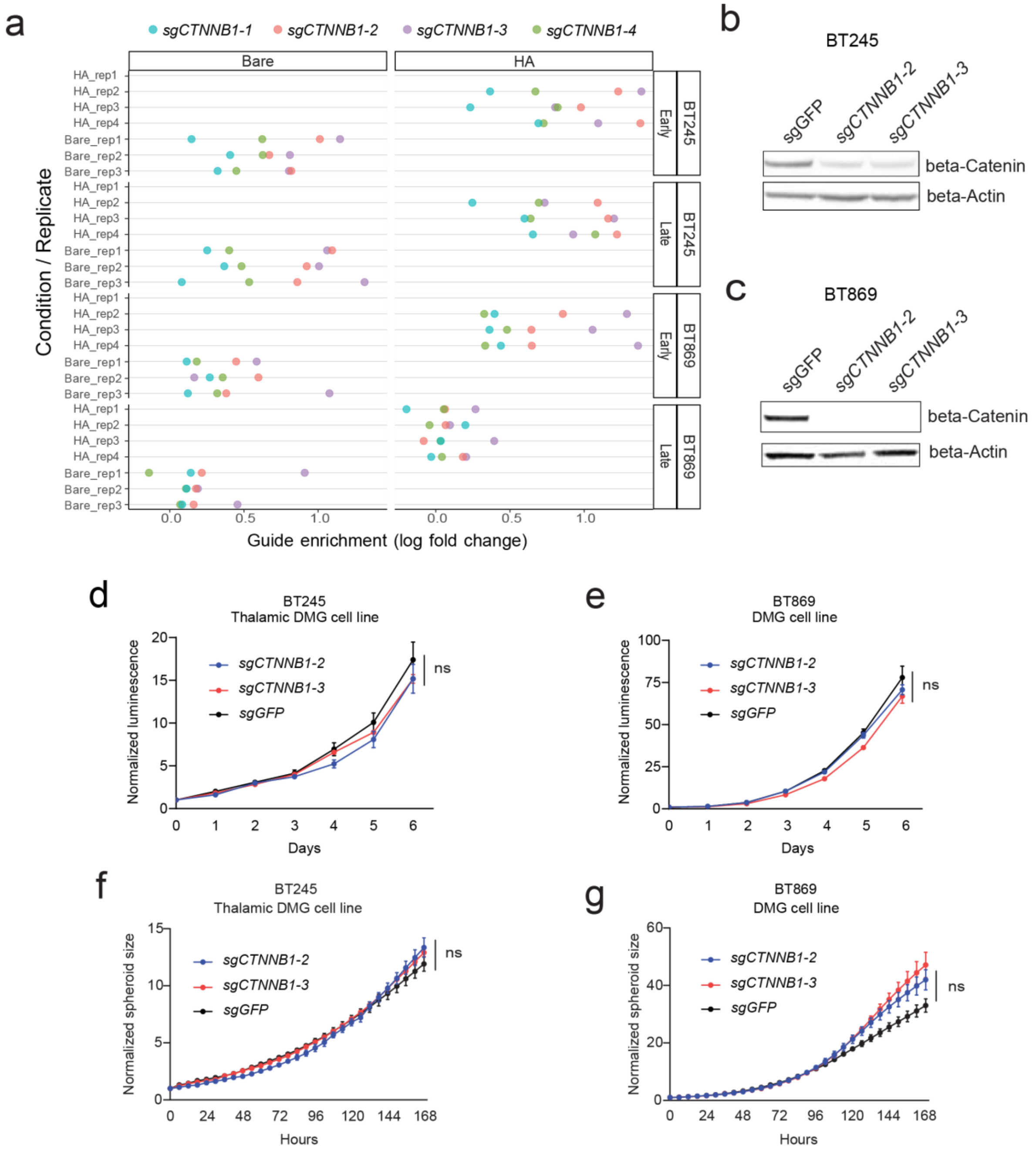


**Supplementary Fig. 5 | *CTNNB1* single guide RNA selection and gene dependency in DMG cell lines. a,** Guide enrichment data of technical replicates in DMG models incubated with liposomes and sorted at early and late timepoints. Western blot validation of β-catenin knockout in (**b**) BT245 and (**c**) BT869 cell lines. Cell proliferation evaluated using (**d**)(**e**) CellTiter-Glo assay and (**f**)(**g**) spheroid size measurement using Incucyte spheroid module. Values indicate mean ± SEM across three replicates. ns $p>0.05$ determined by Dunnett's multiple comparisons test of *sg2CTNNB1* and *sg3CTNNB1* relative to *sgGFP* at 168-hours timepoint.

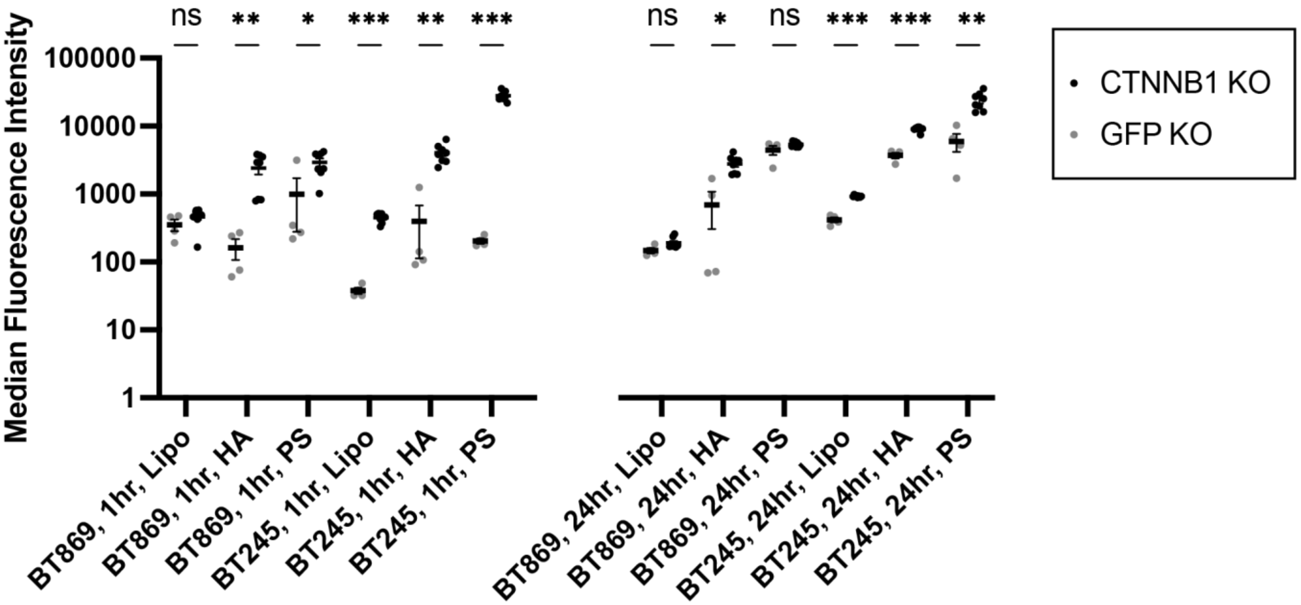


**Supplementary Fig. 6 | Raw median fluorescence intensity data for *CTNNB1* knockout experiments in BT245 and BT869 models for incubation times not shown in Fig. 3D.** Data shown after 1-hour incubation (left) and 24-hour incubation (right) with three different nanoparticles (bare liposome, Lipo; hyaluronic acid-coated liposome, HA; polystyrene, PS). This experiment was performed at a late timepoint (11 days) after infection and selection. Statistical significance assessed using two-tailed Welch's t-test, ** P<0.01, *** P<0.001). Error bar indicates standard error of mean.

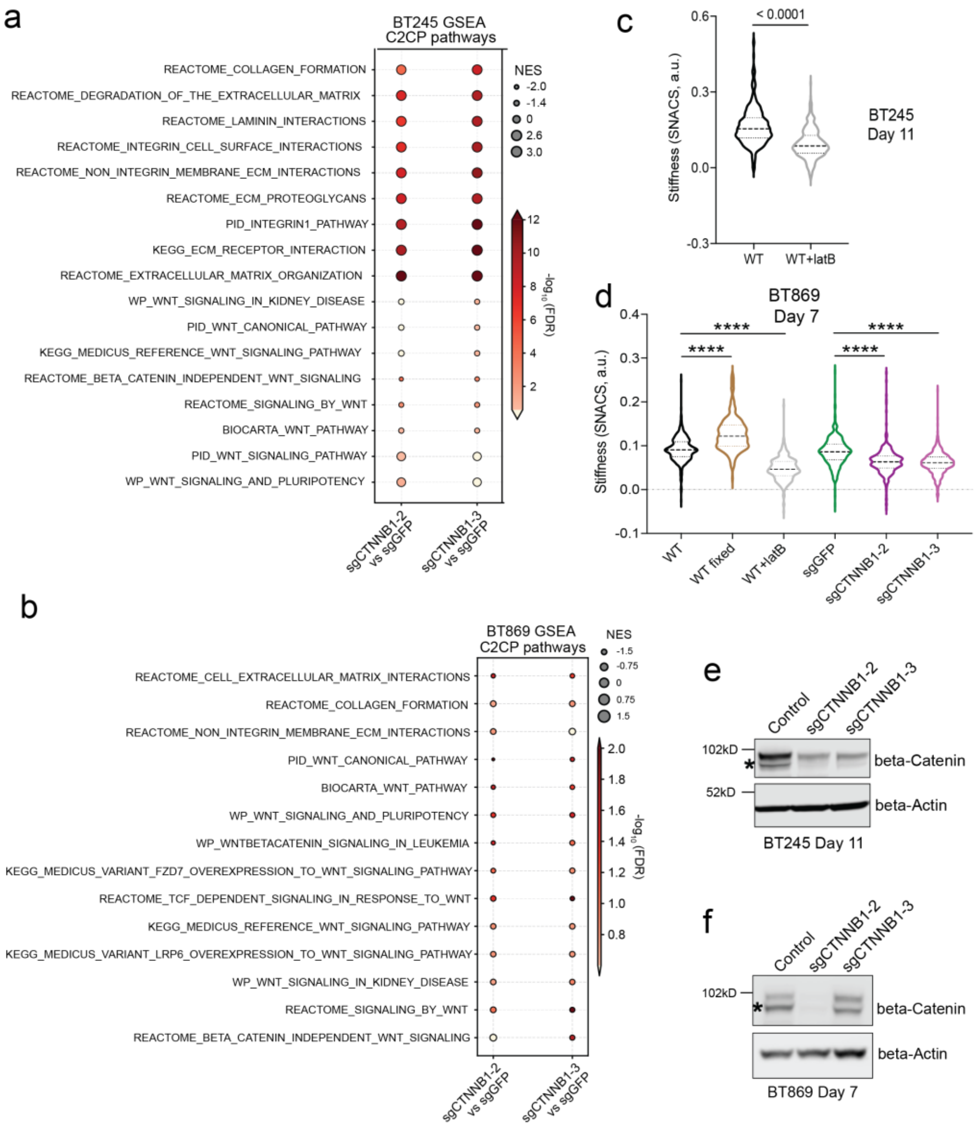


**Supplementary Fig. 7 | *CTNNB1* knockout changes cell membrane mechanics in diffuse midline glioma models.** Dot plot depicting the gene set enrichment analysis (GSEA) results for *CTNNB1* knockout cells using two independent guides in (**a**) BT245 and (**b**) BT869 DMG model. The figure includes all significantly enriched and depleted C2:CP pathways below FDR 0.05., compared to control cell line. The pathways are ranked by NES. **c**-**d**. Violin plots showing normalized cellular stiffness measurements for WT and Latrunculin B treated BT245 cell line model (**c**), control and *CTNNB1* KO BT869 cell lines at day 7 post-selection (**c**), and control and *CTNNB1* KO BT245 cell lines at day 11 post-selection (**d**). Statistical significance was assessed using Welch's t-test, with **** indicating $p < 0.0001$ and ** indicating $p < 0.01$. **e-f**, Western blot analysis showing β-catenin protein levels in control and *CTNNB1* KO DMG cell line models (BT245 and BT869), at different times post-selection (Day 11 and 7).

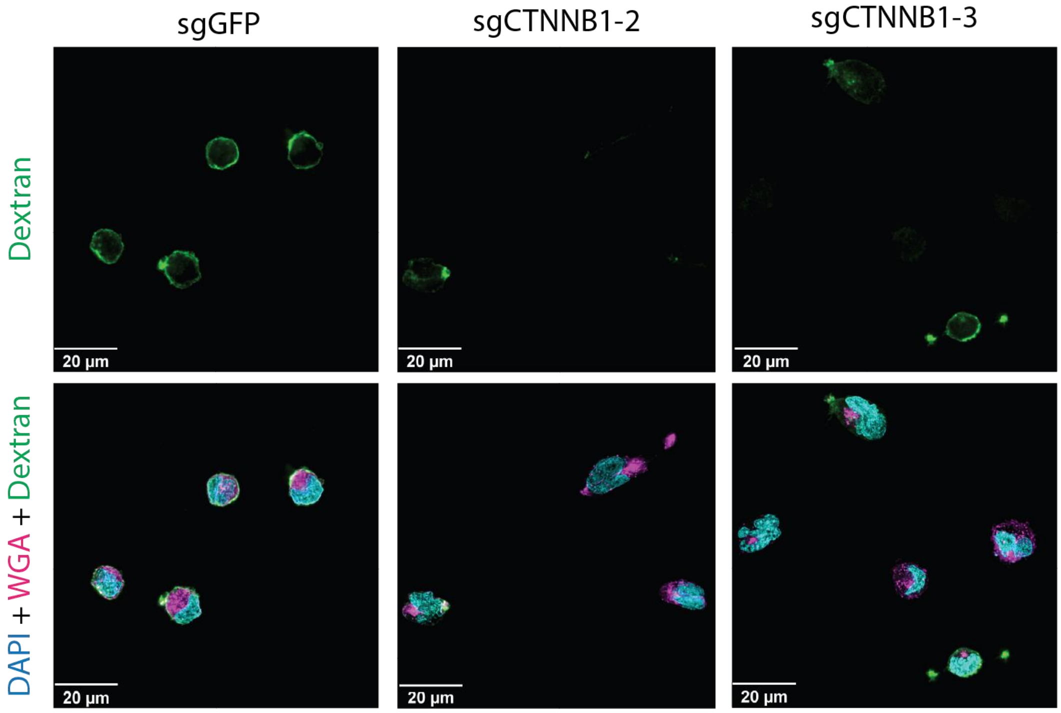


**Supplementary Fig. 8 | Z-projection of middle slices collected from 3D fluorescence microscopy of BT245 sgGFP control and *CTNNB1* knockout cells treated with dextran, a macropinocytosis probe.**

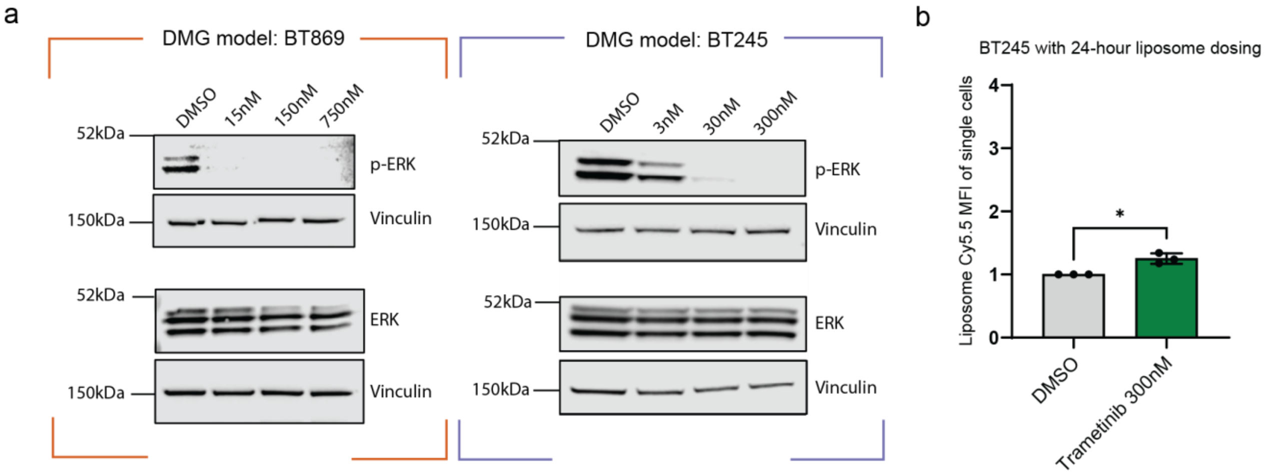


**Supplementary Fig. 9 | MEK inhibitor validation in both DMG models and nanoparticle association in BT245. a**, Western blot evaluation of MEK inhibitor with different concentrations by probing phosphorylated ERK. **b**, Normalized MFI of liposome-association to MAPK inhibitor-treated BT245 measured with flow cytometry. Statistical significance was assessed using Welch's t-test, with * indicating $p < 0.05$.
